# Lipid droplets disrupt mechanosensing in human hepatocytes

**DOI:** 10.1101/2020.03.31.017319

**Authors:** LiKang Chin, Neil D. Theise, Abigail E. Loneker, Paul A. Janmey, Rebecca G. Wells

## Abstract

Hepatocellular carcinoma (HCC) is the fourth leading cause of cancer death in the world. Although most cases occur in stiff, cirrhotic livers, and stiffness is a significant risk factor, HCC can also arise in non-cirrhotic livers in the setting of non-alcoholic fatty liver disease (NAFLD). We hypothesized that lipid droplets in NAFLD might apply mechanical forces to the nucleus, functioning as mechanical stressors akin to stiffness. We investigated the effect of lipid droplets on cellular mechanosensing and found that primary human hepatocytes loaded with the fatty acids oleate and linoleate exhibited decreased stiffness-induced cell spreading and disrupted focal adhesions and stress fibers. The presence of large lipid droplets in hepatocytes resulted in increased nuclear localization of the mechano-sensor Yes-associated protein (YAP). In cirrhotic livers from patients with NAFLD, hepatocytes filled with large lipid droplets showed significantly higher nuclear localization of YAP as compared to cells with small lipid droplets. This work suggests that lipid droplets induce a mechanical signal that disrupts the ability of the hepatocyte to sense its underlying matrix stiffness, and that the presence of lipid droplets can induce intracellular mechanical stresses.

**GRAPHICAL ABSTRACT:** 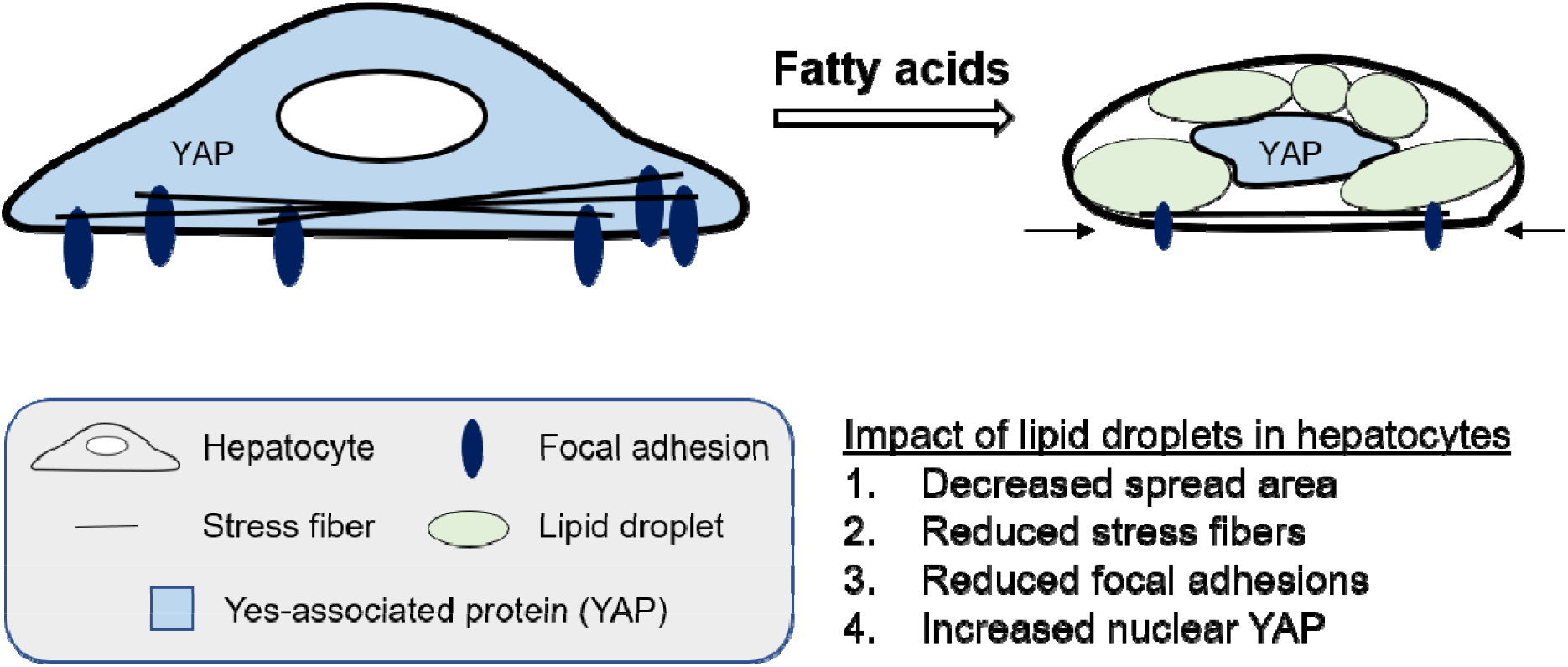

## INTRODUCTION

Hepatocellular carcinoma (HCC) is the sixth most prevalent cancer (6) and the fourth leading cause of cancer mortality worldwide (33). Although the most common risk factor for HCC is currently chronic viral hepatitis, non-alcoholic fatty liver disease (NAFLD) is an increasing cause of HCC and is expected to overtake viral hepatitis in the next few decades (13, 24, 30).

Eighty to ninety percent of HCC cases arise in a stiff, cirrhotic liver (1, 12, 17), and high liver stiffness correlates strongly with an increased risk of HCC (1, 15, 20, 28). Tissue stiffness is a contributing factor to cancer development in many tissues (2, 4, 27), and *in vitro* studies with hepatocytes show that gene expression, motility, and cell stiffness are altered in response to substrate stiffness (31, 32). Hepatocytes cultured on a stiff matrix dedifferentiate, with decreased albumin production and glycogen storage and disruption of the HNF4-α transcriptional network (8).

HCC also occurs in non-cirrhotic and, rarely, normal livers; this is particularly true for HCC in NAFLD, where up to a third of cases occur in non-cirrhotics (12, 17). The defining feature of NAFLD is lipid accumulation in hepatocytes, ranging from small to intermediate to large droplet fat. The largest droplets may be so large that they occupy nearly the entire cytoplasmic space, displacing the deformed nucleus to the periphery. Small and intermediate droplet fat appears as numerous lipid droplets filling the cell with a still centrally-located nucleus. Mechanical measurements of lipid droplets in adipocytes using atomic force microscopy showed that they are stiffer than the cytoplasm (although not the nucleus), and simulations suggest that lipid droplets can induce significant mean shear strains (up to approximately 36%) in the cytoplasm and contribute to an increase in cell stiffness (23). We hypothesized that lipid droplet-associated mechanical signals in hepatocytes disrupt the ability of cells to detect the underlying matrix stiffness, thereby providing a possible pathway for HCC progression even in the absence of significant fibrosis. We evaluated the localization of the mechano-effector Yes-associated protein (YAP) in fatty livers and the effects of the fatty acids oleate (monounsaturated) and linoleate (polyunsaturated) on the actin cytoskeleton, focal adhesions, and localization of YAP in hepatocytes cultured on polyacrylamide gels of variable stiffness.

## MATERIALS AND METHODS

### Human liver collection

De-identified human liver samples from male and female patients aged 25 to 80 years old who underwent liver resection or transplantation were collected in strict accordance with protocols approved by the Institutional Review Boards of the University of Pennsylvania and the Corporal Michael J. Crescenz Veterans Administration Medical Center. Liver tissue was stored in cold Hank’s Balanced Salt Solution (HBSS, Gibco, Gaithersburg, MD) until fixation or rheological testing, which was carried out within 18 hours of harvest.

### Tissue staining and analysis

Cirrhotic liver samples from patients with various etiologies of liver disease were fixed in 10% neutral buffered formalin (Fisher Scientific, Hampton, NH), paraffin embedded, and cut into 5 μm-thick sections. Slides were de-waxed, rehydrated, and stained with hematoxylin and eosin (H&E). One representative H&E-stained section from each patient sample was qualitatively scored by a blinded, board-certified pathologist (NDT) for the extent of cirrhosis according to the proposed Beijing Classification (26). Additionally, NAFLD was determined from clinical history. Samples with clinically intermediate or advanced stage cirrhosis with or without NAFLD were used for further histological and rheological study (n=27 non-NAFLD, n=9 NAFLD).

For Sirius red, sections were stained for 1 h in picro Sirius red (Poly Scientific R&D, Bay Shore, NY), washed in water, and dehydrated in 10% ethanol. For YAP staining, a subset of samples (n=6 non-NAFLD cirrhotic and n=5 NAFLD cirrhotic livers) were subjected to antigen retrieval in 10 mM sodium citrate buffer (pH 6.0) using a pressure cooker, treated with 3% hydrogen peroxide for 20 min at RT to block endogenous peroxidase activity, and permeabilized with 0.1% Triton X-100 (Sigma-Aldrich, St. Louis, MO) in PBS for 10 min at RT. After blocking with 10% normal goat serum (Jackson ImmunoResearch Labs, West Grove, PA) and 0.05% Tween 20 (Bio-Rad, Hercules, CA) in PBS for 1.5 h at RT, slides were incubated with rabbit anti-YAP antibody (1:400, 14074, Cell Signaling Technology, Danvers, MA) overnight at 4°C. The next day, sections were blocked for endogenous avidin and biotin activity and incubated with a biotinylated anti-rabbit IgG secondary antibody (0.5%, Vector Laboratories, Burlingame, CA) for 30 min at RT, followed by Vectastain ABC reagent (Vector) for 30 min at RT. Finally, sections were incubated with DAB (3,3’-diaminobenzidine, Vector) for 3 min and counterstained with Gill’s hematoxylin (Leica, Buffalo Grove, IL) for 30 sec. After staining, sections were cleared in xylene and mounted with Cytoseal mounting medium (Richard-Allan Scientific, San Diego, CA). A small subset of randomly-selected cirrhotic samples (n=5 non-NAFLD, excluding alcoholic cirrhosis, n=6 NAFLD) were embedded in Tissue-Tek Optimal Cutting Temperature (OCT) compound (Sakura, Torrance, CA), cut into 5 μm-thick frozen sections, and stained with Oil Red O (ORO). Sections were fixed in 10% NBF for 10 min, incubated in propylene glycol (Fisher Chemical) for 2 min, and in ORO solution (Poly Scientific R&D) overnight at 4°C. Sections were incubated in 85% propylene glycol and water for 1 min each, then mounted with glycerin jelly (Poly Scientific R&D).

Stained sections were imaged using a Nikon E600 brightfield microscope. At least nine representative spot images were taken, evenly spaced across the surface of the tissue in a grid pattern. Using open-source Fiji software, each image was converted to 8-bit, a pixel threshold was applied (0-120 for ORO, 0-110 for Sirius Red), and the percent area of staining was quantified. For YAP-stained sections, the staining intensity of the nucleus versus cytoplasm was qualitatively compared for each cell, and the percentage of cells with more nuclear than cytoplasmic YAP staining was quantified. NAFLD livers were evaluated for both micro- and macrovesicular steatosis. Microvesicular steatosis was defined as the presence of numerous small lipid droplets and a round nucleus centered within the cell; macrovesicular steatosis was defined as the presence of one large lipid droplet displacing and compressing a nucleus towards the cell membrane.

### Rheometry

Liver samples were cut into 8 mm diameter discs with a stainless-steel punch. The shear storage modulus G’ was measured by applying a low oscillatory shear strain of 2% at a frequency of 1 rad/s using a strain-controlled rotational RFS3 rheometer (TA Instruments, New Castle, DE, USA) fitted with parallel plates. Fibrin glue prepared by mixing 5 μl of 28 mg/ml salmon fibrinogen with 5 μl of 10 units of salmon thrombin, both purified as described (29), was used to adhere samples to the rheometer plates to avoid slippage during shear deformation. The top plate was lowered until contact was made as determined by the application of 1.7 g of normal force. Samples were kept hydrated with PBS during experiments.

### Glycosaminoglycan measurement

Sulfated glycosaminoglycan (sGAG) concentration was determined from a subset of randomly-selected samples using the Blyscan reagent according to the manufacturer’s instructions (Biocolor, UK). Small chunks of cirrhotic liver (n=20 non-NAFLD, n=5 NAFLD, up to 50 mg) were placed in 1 ml of digestion buffer (0.2 M sodium phosphate buffer, pH 6.4, 0.1 M sodium acetate, 10 mM EDTA, 0.08% cysteine HCl (Sigma) and 0.02% papain (Worthington, Lakewood, NJ)) and incubated at 65°C for 18 h. The solution was centrifuged at 10,000 x g for 10 min. One hundred μl of supernatant was combined with 1 ml of the Blyscan dye reagent, incubated for 30 min at room temperature with shaking, and centrifuged at 12,000 x g for 10 min. The resulting pellets were dried and dissolved in 0.61 ml of the provided dissociation reagent. Absorbance was read from triplicate wells at 656 nm using a Tecan Infinite 200 Pro spectrophotometer (Morrisville, NC), and concentration was determined by comparison to a calibration curve constructed from manufacturer-provided GAG standards.

### Hyaluronan measurement

Hyaluronan (HA) concentration was determined using the Purple-Jelley assay, according to the manufacturer’s instructions (Biocolor). Small chunks of cirrhotic liver (n=20 non-NAFLD, n=5 NAFLD, up to 50 mg) were digested with 400 μl of 250 μg/ml of proteinase K (Sigma) in PBS at 55°C for 18 h and centrifuged at 12,000 rpm for 10 min to remove undigested tissue residue. GAGs in the supernatant fraction were precipitated by adding 1 ml of the provided GAG precipitation reagent followed by centrifugation at 12,000 rpm for 10 min. The resulting GAG pellet was resuspended in 360 μl of water, 40 μl of 23.3% w/v NaCl, and 95 μl of 2% w/v cetylpyridinium chloride, centrifuged at 12,000 rpm for 10 min, and the supernatant was collected. The GAG precipitation-resuspension step was repeated once more, followed by one additional GAG precipitation step. Five hundred μl of 98% ethanol was added to the pellet, followed by centrifugation at 12,000 rpm for 5 min. The remaining pellet was isolated, solubilized in 100 μl of water, and 1 ml of dye reagent was added. Absorbance was read from triplicate wells at 656 nm using a Tecan Infinite 200 Pro spectrophotometer (Morrisville, NC), and concentration was determined by comparison to a calibration curve constructed from manufacturer-provided HA standards.

### Polyacrylamide gel preparation

Polyacrylamide (PAA) gels with storage modulus values of 500 and 10 kPa, representing the stiffness of normal and cirrhotic livers, respectively, were made as previously described (34). Gels and glass substrates activated with sulfo-SANPAH (Thermo Fisher Scientific, Waltham, MA) were incubated with 0.1 mg/ml rat tail collagen type I (Corning, Edison, NJ) for 2 h to coat the top surface with adhesive extracellular matrix ligands. For Huh7 cell culture, glass coverslips were left uncoated.

### Cell culture

Cryo-preserved primary human hepatocytes from single donors (PHH; BD Gentest, Tewksbury, MA; Lonza, Walkersville, MD; Thermo Fisher Scientific) and the human HCC cell line HuH7 (ATCC, Manassas, VA) were used for *in vitro* experiments. PHH were thawed according to the supplier’s instructions using Cryopreserved Hepatocyte Recovery Medium (CM7000, Thermo Fisher Scientific), then resuspended in Williams Medium E (Sigma-Aldrich) with plating supplements (CM3000, Thermo Fisher Scientific) and plated onto collagen-coated PAA gels, glass, or 96-well plates at a density of 50,000 cells per cm^2^. HuH7 cells were cultured in DMEM (Corning) supplemented with 10% FBS (Gemini Bio-Products, West Sacramento, CA) and 1% penicillin-streptomycin (Corning) and seeded onto collagen-coated PAA gels or uncoated glass at a density of 18,750 cells per cm^2^. For positive control cells for apoptosis, PHH were treated with 5 μM staurosporine (STS) for 20 h one day after seeding.

### Fatty acid treatment

One day after seeding, PHH and HuH7 cells were serum starved overnight in either serum-free Williams Medium E with maintenance supplements (CM4000, Thermo Fisher Scientific) or DMEM supplemented with 0.2% free fatty acid-free bovine serum albumin (FFA-free BSA, Sigma-Aldrich) and 1% penicillin-streptomycin, respectively. Cells were then incubated for 48 h in the presence or absence of 400 μM sodium oleate or linoleic acid sodium salt (≥98%, Sigma-Aldrich) in DMEM supplemented with 0.5% BSA and 1% penicillin-streptomycin. To solubilize the fatty acids, 20 mM fatty acid solutions were prepared in 0.01 M NaOH and incubated for 30 min at 70°C, then diluted to 4 mM with 5% FFA-free BSA in PBS and incubated at 55°C for 10 min to conjugate the fatty acids to BSA. The fatty acid solutions were then mixed 1:9 with serum-free DMEM with 1% penicillin-streptomycin to obtain a 400 μM fatty acid, 0.5% BSA solution in DMEM (7, 22). To evaluate the effect of lipid droplet size on YAP signal, cells were treated with 400 μM oleate and 100 nM insulin (Sigma-Aldrich) for 48 h to inhibit lipolysis and generate large lipid droplets.

### Cytotoxicity assay

Cytotoxicity was measured in triplicate samples using the lactate dehydrogenase (LDH) cytotoxicity assay (Thermo Fisher Scientific) as directed by the manufacturer. Briefly, PHH were seeded at 50,000 cells per cm^2^ onto collagen-coated tissue culture plastic and treated with or without fatty acids. After 48 h, 50 μl of culture medium from each sample was mixed with 50 μl of reaction mixture, incubated for 30 min at RT, and then the reaction was terminated with the addition of 50 μl of stop solution. Absorbance was measured at 490 nm with absorbance at 680 nm subtracted as background using a Tecan Infinite 200 Pro spectrophotometer. For comparison, maximum LDH release was measured from cells treated with 10 μl lysis buffer.

### Cell staining, microscopy, and analysis

To assess cell shape factors (cell area, circularity, and solidity), unfixed, unstained cells were imaged at 10X using a Leica DMi1 tissue culture microscope in the same session and under the same conditions. For immunofluorescence staining, cells were fixed in 4% paraformaldehyde (PFA, Thermo Fisher Scientific) for 10 min and permeabilized with 0.1% Triton X-100 in PBS for 15 min. For vinculin, samples were simultaneously fixed and permeabilized with microtubule stabilizing buffer (0.1 M PIPES, pH 6.75 (Honeywell Fluka. Muskegon, MI), 1 mM EGTA (Calbiochem, San Diego, CA), 1 mM MgSO_4_ (Sigma-Aldrich), 4% polyethylene glycol (United States Pharmacopeia, North Bethesda, MD), 1% Triton X-100, and 2% PFA) for 10 min at 37°C (3). Samples were blocked with 5% normal goat serum (Jackson ImmunoResearch Labs, West Grove, PA) in PBS for 1 h. To stain neutral lipids, cells were incubated with BODIPY (1:1000, Thermo Fisher Scientific) for 1.5 h at 37°C. For immunostaining, primary antibodies (rabbit anti-YAP, 1:100; rabbit anti-vinculin, 1:100, ab129002, Abcam, Cambridge, UK; rabbit anti-cleaved caspase-3, 1:400, 9446, Cell Signaling) were applied overnight at 4°C. Cells were then incubated in secondary antibody (fluorophore-conjugated anti-rabbit IgG, 1:600, Jackson ImmunoResearch Labs or poly-HRP-conjugated anti-rabbit IgG, undiluted, Invitrogen, Carlsbad, CA) for 1 h at RT. To boost the YAP and vinculin signals, cells were incubated in Tyramide Working Solution for 3.5 min, followed by Reaction Stop Reagent (both from Tyramide SuperBoost Kit, Invitrogen). Where applicable, actin was stained with phalloidin (1:50, Invitrogen) for 20 min at RT, and nuclei with Hoescht (1:10,000, Thermo Fisher Scientific) for 10 min at RT. Samples were then mounted with aqueous mounting medium (KPL, Gaithersburg, MD) and 40X images were taken using a Zeiss LSM 710 laser scanning confocal during the same session and under the same conditions.

To assess cell senescence, BSA control and fatty acid treated-cells were stained for β-galactosidase. Cells treated with 300 μM H_2_O_2_ for eight days served as positive control. After fixation in a 0.2% glutaraldehyde (Fisher Scientific), 1.8% PFA solution in PBS for 5 min at RT, cells were stained overnight at RT with 1 mg/ml X-gal in a 5 mM potassium ferricyanide, 5 mM potassium ferrocyanide, 2 mM MgCl_2_, 150 mM NaCl, 40 mM citric acid/sodium phosphate buffer (all from Sigma). Cells were imaged at 10X using a Leica DMi1 tissue culture microscope during the same session and under the same conditions.

Cell morphology, mean intensity, and integrated density were analyzed by manually tracing images of single, non-touching cells using Fiji. For YAP-stained cells, YAP mean intensity of both the nucleus and the entire cytosol was measured, and the ratio of nuclear to cytosolic YAP was calculated. For vinculin-stained cells, signal intensity was optimized for each image to visualize the presence or absence of actin stress fibers and vinculin patches, defined as punctate staining located at the end of a stress fiber. The total area of vinculin staining per cell, including patches and cytosolic vinculin, was quantified from images taken with identical settings.

To determine the effect of lipid droplet size on YAP signal, cells treated with either oleate alone or both oleate and insulin were co-stained for neutral lipids and YAP. Lipid droplet size for each cell was qualitatively assessed in a systematic way and binned as either small, medium, or large. The nuclear to cytosolic YAP ratio and nuclear circularity were determined.

### Statistical analysis

Unpaired t tests were performed to examine the effects of NAFLD on G’, Sirius Red and ORO staining, as well as sGAG and HA quantification. G’-Sirius Red, G’-ORO, G’-sGAG, and G’-HA data were fit with a power law function, and correlation tests were run. Two-way ANOVA was performed to examine the effects of substrate stiffness and fatty acids on lipid storage, cell/nuclear morphology, cytoskeletal structures (actin stress fibers and focal adhesions), as well as YAP localization and intensity. One-way ANOVA was used to test the statistical significance of fatty acids on cytotoxicity and caspase3 staining intensity and also the effect of steatosis on nuclear YAP in human samples. When appropriate, multiple comparisons were performed with Tukey’s post-hoc test. For all statistical analyses, a p-value of ≤ 0.05 was considered significant. Results are expressed as mean ± standard error of the mean (SEM).

## RESULTS

### Liver stiffness, collagen, sGAG, and HA were similar in NAFLD and non-NAFLD cirrhosis

Human livers with intermediate (characterized by focal or frequent fibrous septae) or advanced (focal or diffuse nodularity) cirrhosis per the proposed Beijing Classification system underwent *ex vivo* measurements of liver stiffness (shear storage modulus G’) using parallel plate rheometry (Fig. 1A). No statistically significant difference was found between the stiffness values for non-NAFLD cirrhotic livers (932.6 ± 51.1 Pa) and NAFLD cirrhotic livers (856.9 ± 123.7 Pa, Fig. 1B), although the stiffness of the non-NAFLD cirrhotic livers was less variable than NAFLD cirrhotic livers, with coefficients of variance of 26.5% and 41.0%, respectively. Similarly, there was no statistically significant difference between the two groups for Sirius Red staining (Fig. 1C) or sGAG (Fig. 2A) or HA content (Fig. 2B). Cirrhotic livers from patients with a documented history of NAFLD had more ORO staining than non-NAFLD livers (p=0.027, Fig. 1D). Liver stiffness correlated, but non-linearly, with Sirius Red staining (Fig. 1E), and there was no correlation with lipid (Fig. 1F), sGAG (Fig. 2C), or HA content (Fig 2D).

**Fig. 1.**
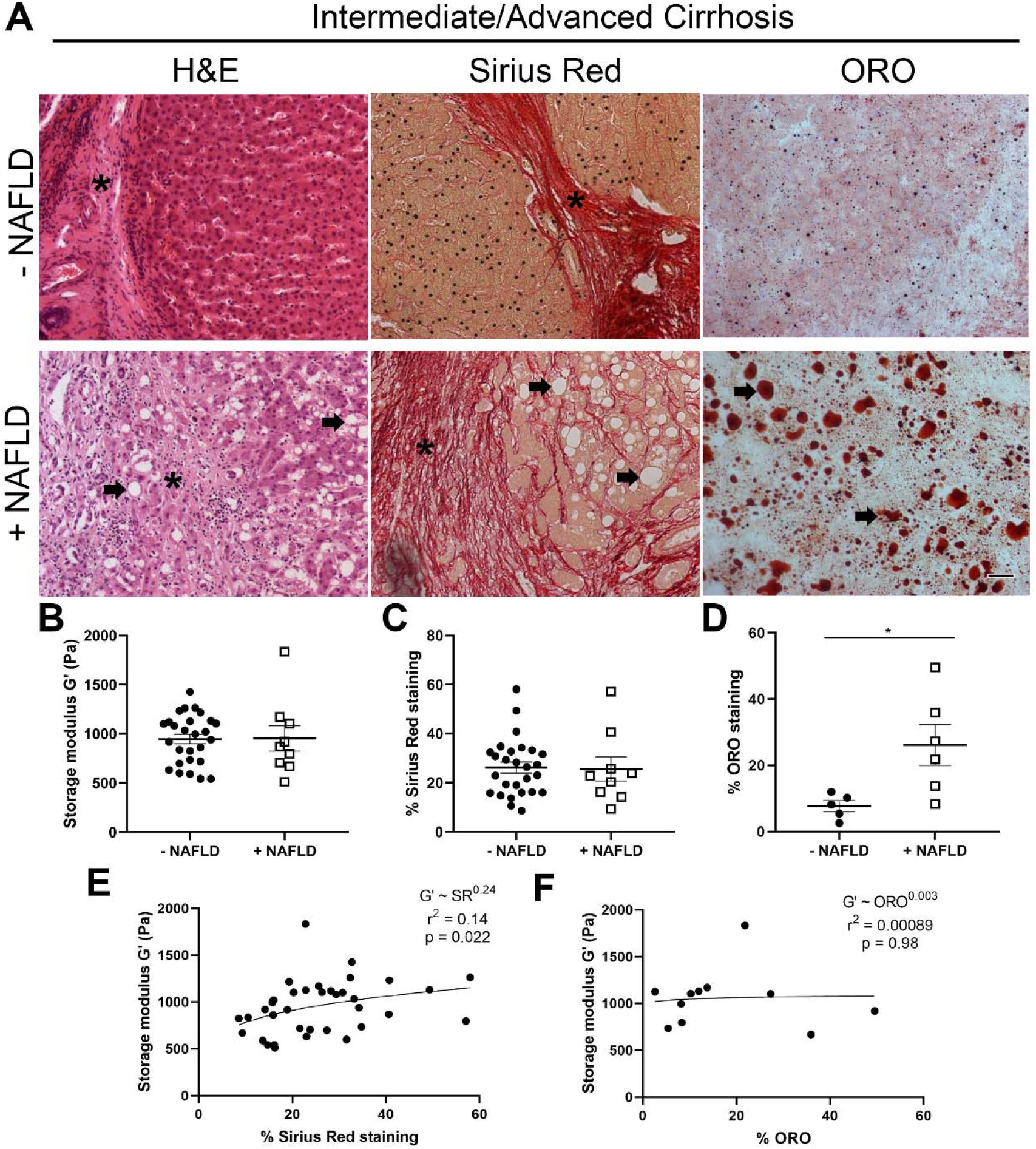
NAFLD livers have more lipid, but similar stiffness and collagen as non-NAFLD cirrhotic livers. **A)** Representative images of H&E-, Sirius Red-, and ORO-stained human livers staged as intermediate/advanced cirrhosis, from patients with or without a documented clinical history of NAFLD. Livers exhibited nodules surrounded by fibrous septa (asterisks), and those with NAFLD also had large lipid droplets (arrows). **B)** Shear storage modulus G’ of human livers, from patients with or without a history of NAFLD. **C)** Collagen content, as determined by percentage of Sirius red staining, in cirrhotic livers was similar with or without NAFLD. **D)** NAFLD livers had more ORO staining than non-NAFLD livers (p=0.027). Scatter plots of G’ versus **E)** % Sirius Red staining showing nonlinear correlation (power law fit, p=0.022) and **F)** % ORO staining showing no correlation (power law fit, p=0.8). Scale bar, 50 μm. Error bars are SEM. G’, Sirius Red, n=27 non-NAFLD, 9 NAFLD. ORO, n=5 non-NAFLD, 6 NAFLD.

**Fig. 2.**
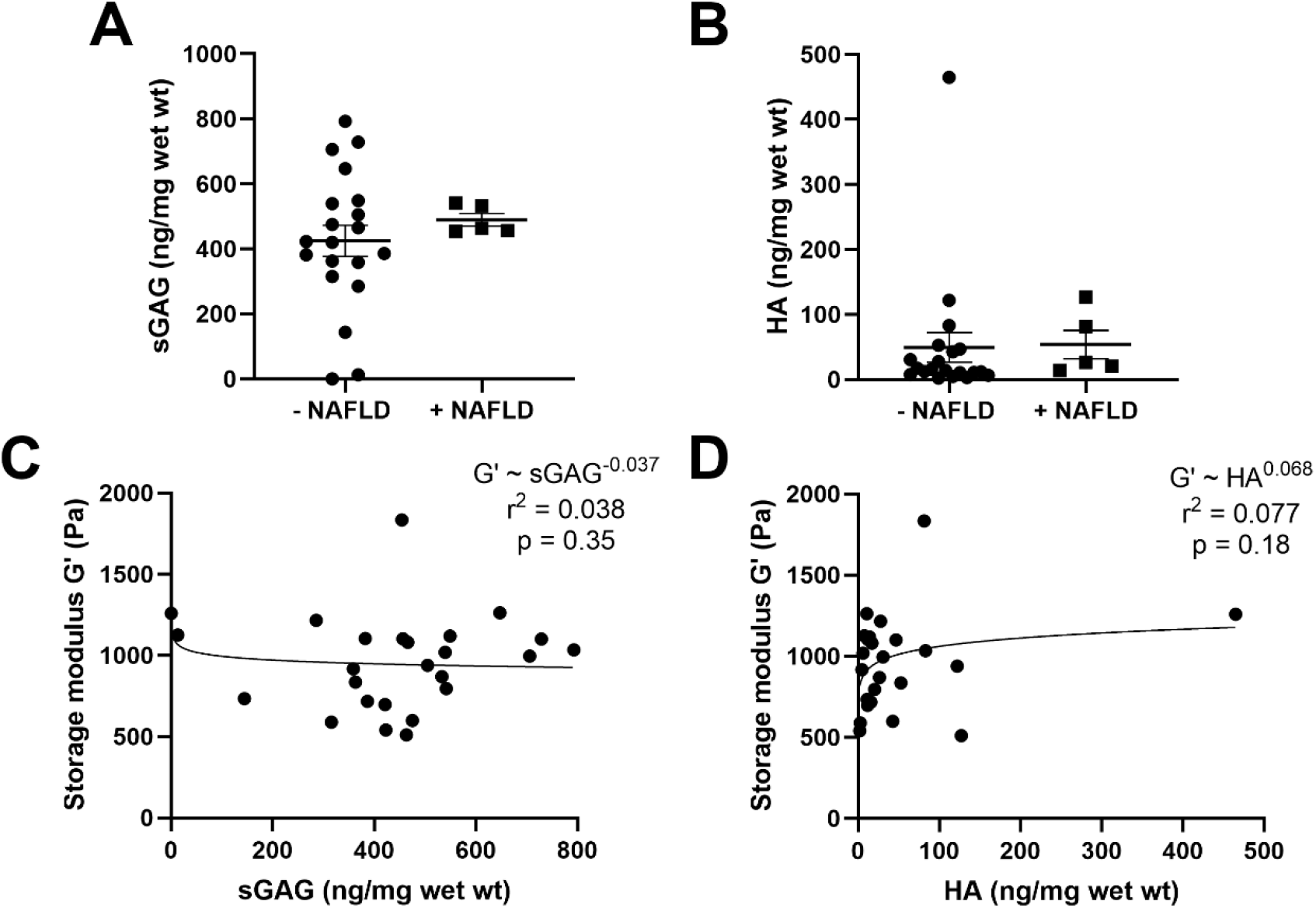
NAFLD cirrhotic livers have the same amounts of sGAG and HA as non-NAFLD cirrhotic livers. **A)** sGAG and **B)** HA quantification for cirrhotic livers, with and without NAFLD. Scatter plots of G’ versus **C)** sGAG showing no correlation (power law fit, p=0.35) and **D)** HA showing no correlation (power law fit, p=0.18). Error bars are SEM. n=20 non-NAFLD, 5 NAFLD.

### Nuclear YAP localization remains unchanged in microvesicular NAFLD and increases in macrovesicular NAFLD

To determine YAP localization in human clinical samples, a set of cirrhotic livers, with or without NAFLD, was stained for YAP. Hepatocytes in NAFLD livers exhibited both micro- and macrovesicular steatosis, and both YAP+ and YAP-nuclei (Fig. 3A). The percentage of individual cells with YAP+ nuclei was quantified for each type of steatosis. Hepatocytes from both control livers (cirrhotic, non-steatotic) and those with microvesicular steatosis had a similar percentage of YAP+ nuclei (approximately 65%, Fig. 3B). Interestingly, despite there being no change in liver stiffness (Fig. 1B), nearly all hepatocytes with macrovesicular steatosis (filled with one or few large lipid droplets displacing and deforming the nucleus) had YAP+ nuclei. In light of these results, we explored the effect of lipid droplets on cell mechanical behavior.

**Fig. 3.**
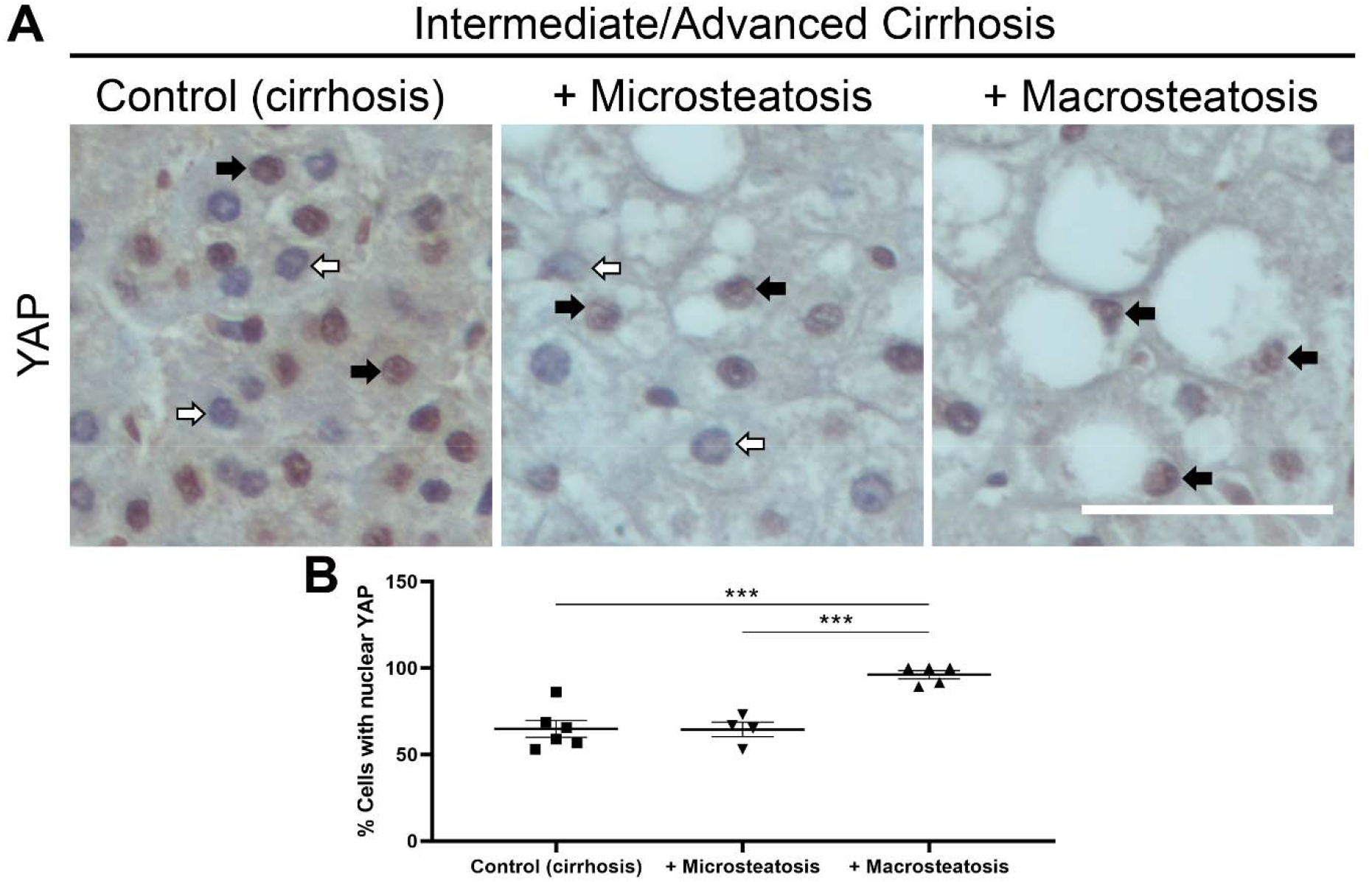
Nuclear YAP localization increases in macrovesicular steatosis. **A)** Representative images of human livers with intermediate or advanced cirrhosis, with or without NAFLD, stained for YAP. Individual cells from five livers were categorized as having micro- or macrovesicular steatosis. Hepatocytes in cirrhotic livers and those with microvesicular steatosis exhibited both YAP+ (closed arrows) and YAP-(open arrows) nuclei. Hepatocytes with macrovesicular steatosis exhibited deformed nuclei that were almost exclusively YAP+ (closed arrows). **B)** Hepatocytes with macrovesicular steatosis had significantly more nuclear YAP than cells with microvesicular steatosis or cells in non-steatotic cirrhotic livers. Scale bar, 50 μm. *** p≤0.001. Error bars are SEM. Control (cirrhosis), n=116-340 cells from 6 livers; Microsteatosis, n=13-104 cells from 4 NAFLD livers; Macrosteatosis, n=30-164 cells from 5 NAFLD livers. One NAFLD liver had fewer than 10 cells with microsteatosis in the fields examined and was dominated by macrosteatosis; it was excluded from the microsteatosis group.

### Fatty acid treatment did not cause cytotoxicity or induce senescence or apoptosis

To determine the impact of lipid droplets on cell mechanics, we studied fatty acid (oleate and linoleate)-loaded PHH and HuH7 cells. First, we identified a dose of fatty acids (400 μM) that resulted in lipid-laden cells without significant toxicity and is also representative of the fatty acid serum concentration of obese patients (18). The effects of fatty acid treatment on hepatocyte cytotoxicity, senescence, and apoptosis were assessed in PHH seeded onto glass and treated with fatty acids for 48 h. As assayed by LDH release, which is a direct measure of cytotoxicity, treatment with either fatty acid did not cause increased cytotoxicity compared to BSA controls (Supplemental Fig. S1A, supplemental material for this article is available online at https://figshare.com/s/2389f50a30c933a3de66). To assess cellular senescence, fatty acid-treated PHH seeded onto glass were stained for β-galactosidase; they demonstrated minimal staining compared to positive control cells (treated with 300 μM H_2_O_2_ for 7 d, Supplemental Fig. S1B). Fatty acid-treated cells stained for activated caspase3, a marker of apoptosis, showed no increase in staining intensity when compared to BSA-treated cells (Supplemental Fig. S1C, D).

### Oleate-treated cells stored more lipid than cells treated with linoleate, and lipid storage increased on soft gels

To study the impact of lipid uptake in combination with environmental stiffness, we seeded PHH and HuH7 cells onto soft (500 Pa) or stiff (10k Pa) PAA gels or glass and subsequently treated them with medium supplemented with either 0.5% BSA alone (control) or with BSA plus 400 μM oleate or linoleate for 48 h. Treatment with either fatty acid resulted in the presence of numerous small lipid droplets. For both PHH and HuH7, oleate-treated cells stored more total lipid and had significantly greater lipid density than those treated with BSA alone or linoleate (Fig. 4A, B), as quantified by mean intensity (Fig. 4C, D) and integrated density (Fig. 4E, F) after BODIPY staining. The amount of oleate storage was stiffness-dependent, and treated cells on soft gels stored significantly more lipid than those on glass. Oleate-treated HuH7 cells had significantly less lipid uptake than their PHH counterparts, regardless of stiffness (p≤0.0001, not shown graphically in Fig. 4E, F).

**Fig. 4.**
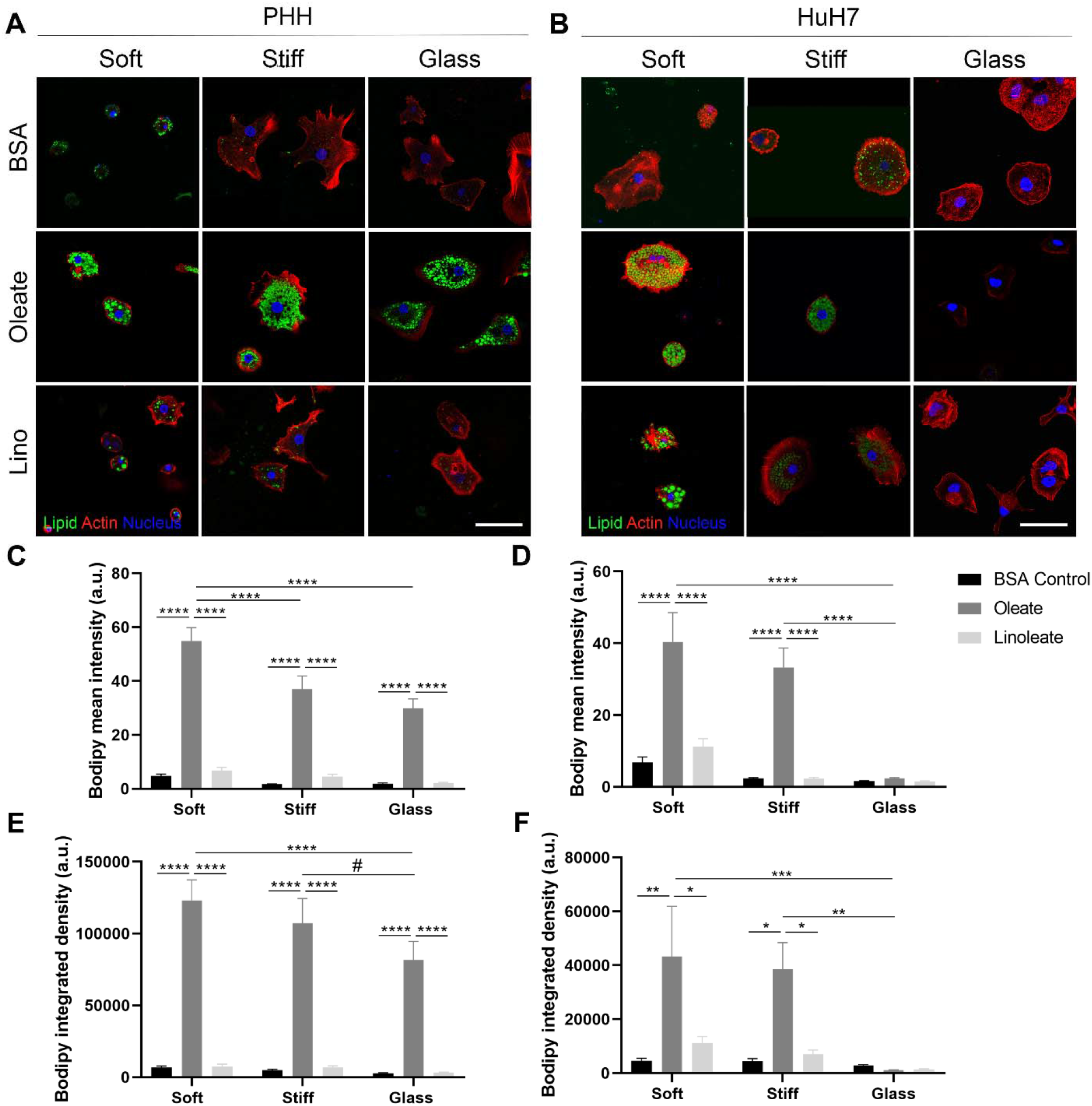
Oleate-treated cells store more lipid than cells treated with linoleate, and lipid storage decreases with stiffness. Representative confocal images of **A)** PHH and **B)** HuH7 cells seeded on 500 Pa (soft) or 10 kPa (stiff) PAA-collagen gels, or glass (collagen-coated for PHH), treated with 400 μM oleate or linoleate for 48 hours, and stained for neutral lipids (green), actin (red), and nuclei (blue). **C, D)** BODIPY mean intensity (a measure of lipid density) and **E, F)** integrated density (a measure of total lipid) were quantified for PHH and HuH7. Oleate-treated HuH7 cells had significantly less lipid density on soft gels and glass (C, D, p≤0.01, not graphically represented) and less total lipid on all substrates (E, F, p≤0.0001, not graphically represented) than their normal PHH counterparts. To better visualize lipid droplets in BSA and linoleate (only PHH) groups, gain was increased all to the same setting; gain was increased in BSA-treated PHH to better see actin. All quantification was performed on images acquired with the same settings. Scale bar, 50 μm. *p≤0.05, ** p≤0.01, ***p≤0.001, ****p≤0.0001. PHH, n=30 total cells per condition from three independent experiments. HuH7, n=12-25 total cells per condition from two independent experiments. Error bars are ± SEM.

### Stiffness-dependent cell spreading decreased in fatty-acid loaded primary human hepatocytes and HuH7 cells

Not surprisingly, both PHH and HuH7 cell spreading was stiffness-dependent, with significantly higher areas on stiff gels and glass than on soft gels (Figs. 5A, B, D, Supplemental Fig. S2). Although lipid loading significantly decreased the spread area of both PHH and HuH7 cells on stiff substrates as compared to BSA controls, PHH, but not HuH7 cells, continued to show some degree of stiffness-dependent spreading (Fig. 5B, D). The shape of PHH was generally unchanged by fatty acid treatment (Fig. 5A, C, F), but HuH7 cells – which typically feature a star-shaped morphology on PAA gels – became more circular and more solid (defined as area divided by convex hull area, ImageJ) after fatty acid treatment (Fig. 5E, G, Supplemental Fig. S2).

**Fig. 5.**
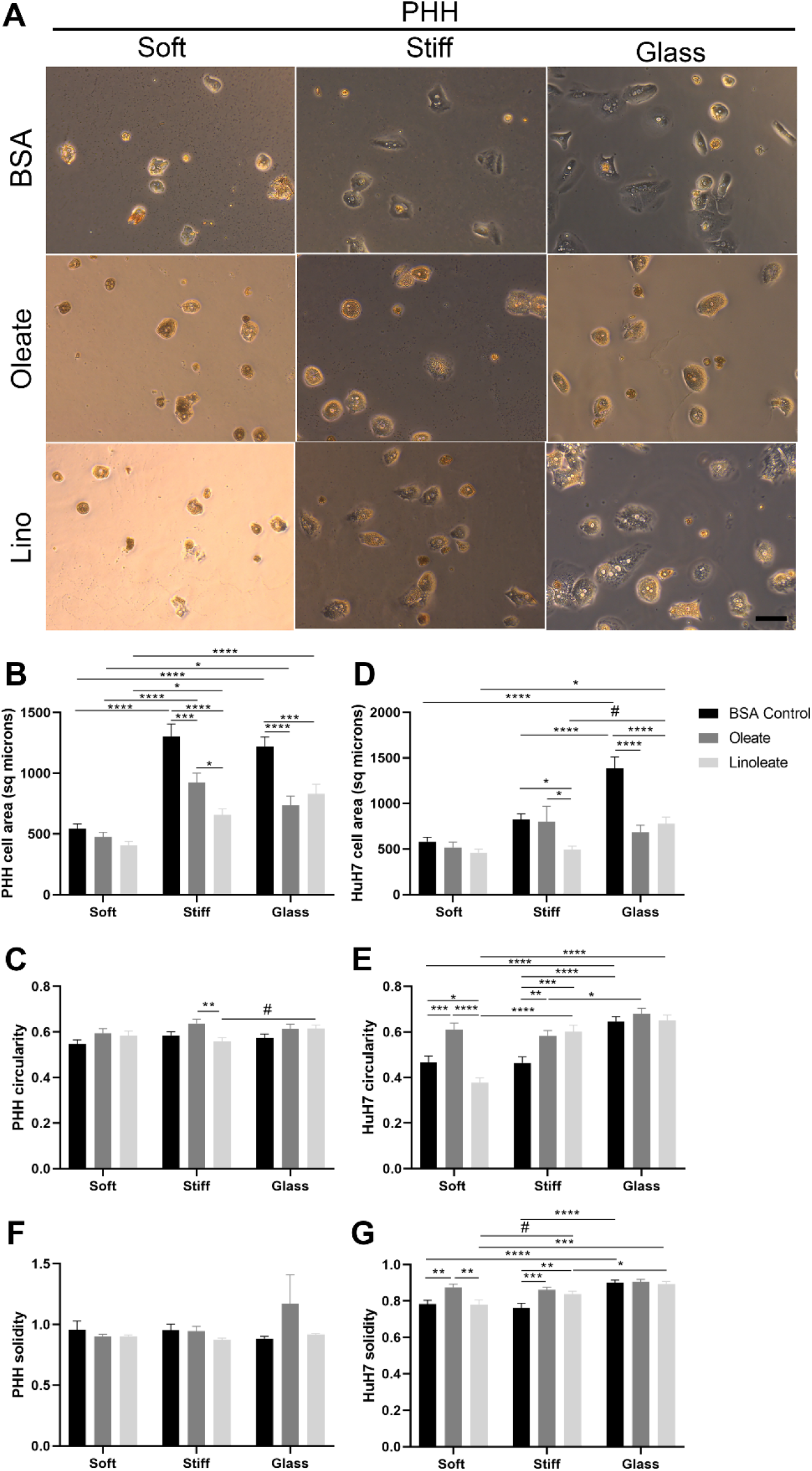
Stiffness-dependent cell spreading decreases in fatty acid-loaded PHH. **A)** Representative live-cell brightfield images of PHH seeded on 500 Pa (soft) and 10k Pa (stiff) PAA-collagen gels, or collagen-coated glass after 48 h of treatment with 400 μM oleate or linoleate. **B, D)** Cell area, **C, E)** circularity, and **F, G)** solidity were quantified for both PHH and HuH7 cells (images shown in Supplemental Fig. S2). Scale bar, 50 μm. *p≤0.05, ** p≤0.01, ***p≤0.001, ****p≤0.0001, #0.05<p≤0.10. n=60 total cells per condition from three independent experiments. Error bars are ± SEM.

### Fatty acid treatment led to disrupted stress fibers and focal adhesions

To visualize stress fibers and focal adhesions, fatty acid-treated and BSA-treated control PHH (Fig. 6A, Supplemental Fig. S3A) and HuH7 cells (Supplemental Fig. S3B, C) on soft and stiff PAA gels and glass were stained for vinculin and actin. The presence of stress fibers and focal adhesions – defined as punctate patches of vinculin staining at the end of stress fibers – was determined for each imaged cell, and total vinculin staining area per cell was quantified. Regardless of substrate stiffness, most BSA-treated PHH exhibited stress fibers (Fig. 6B) and vinculin patches (Fig. 6C,D). These were reduced in lipid-loaded cells, particularly on glass. Fatty acid-treated HuH7 cells showed decreased focal adhesions and vinculin staining on stiff substrates (Supplemental Fig. S3B-F).

**Fig. 6.**
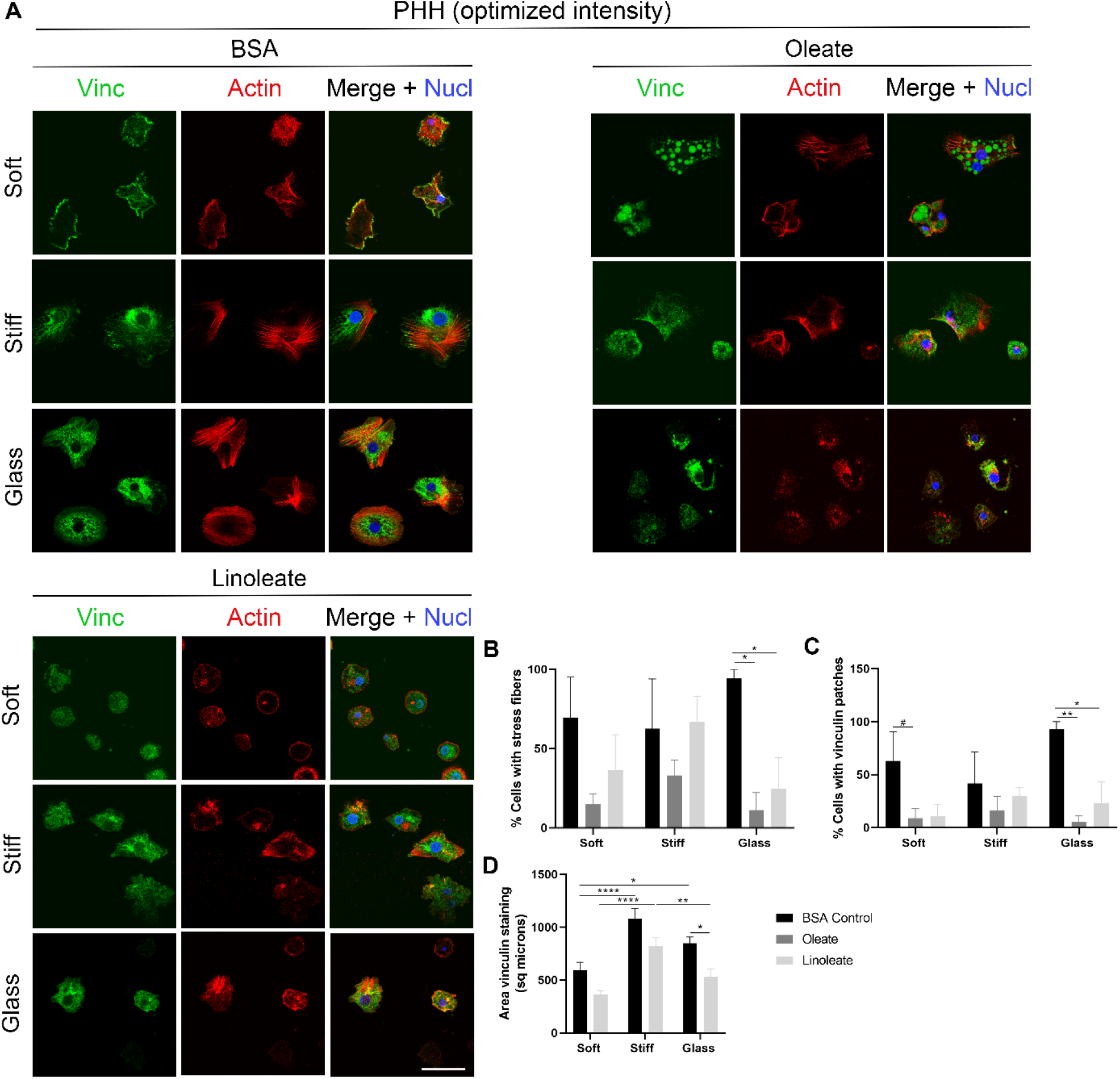
Fatty acid treatment disrupts stress fibers and focal adhesions in PHH, particularly on glass. **A)** Representative confocal images of PHH seeded on 500 Pa (soft) or 10k Pa (stiff) PAA-collagen gels, or collagen-coated glass, treated with 400 μM oleate or linoleate for 48 h, and stained for vinculin (green), actin (red), and nuclei (blue). Quantification is shown for the percentage of cells that exhibited **B)** stress fibers, **C)** vinculin patches, as defined by punctate staining located at the end of stress fibers, and **D)** total cell area of vinculin staining (except for oleate-treated cells due to non-specific staining of the lipid droplets). Intensity was optimized for each image to best visualize the vinculin and actin staining (B, C). Quantification of vinculin staining (D) was from the original images, all taken at the same settings, as shown in Supplemental Fig. S3A. Scale bar, 50 μm. *p≤0.05, ** p≤0.01, ***p≤0.001, ****p≤0.0001, #0.05<p≤0.100. Percent stress fibers or vinculin, n=3 independent experiments. Vinculin area, n=32-63 total cells per condition from three independent experiments. Error bars are ± SEM.

### Cells with large lipid droplets had increased nuclear YAP

To determine whether mechanosensing was altered in lipid-laden cells (which had small lipid droplets using our standard protocol), we stained for YAP, which typically translocates from the cytoplasm to the nucleus with mechanical stress such as that induced by increased substrate stiffness. PHH (Fig. 7A) and HuH7 cells (Supplemental Fig. S4A) showed no stiffness-dependent change in YAP nuclear versus cytoplasmic localization (Fig. 7B, Supplemental Fig. S4B), but increased YAP intensity on glass (Fig. 7C, Supplemental Fig. S4C). Comparison to BSA controls showed that neither YAP localization nor intensity was altered by the presence of lipid droplets.

**Fig. 7.**
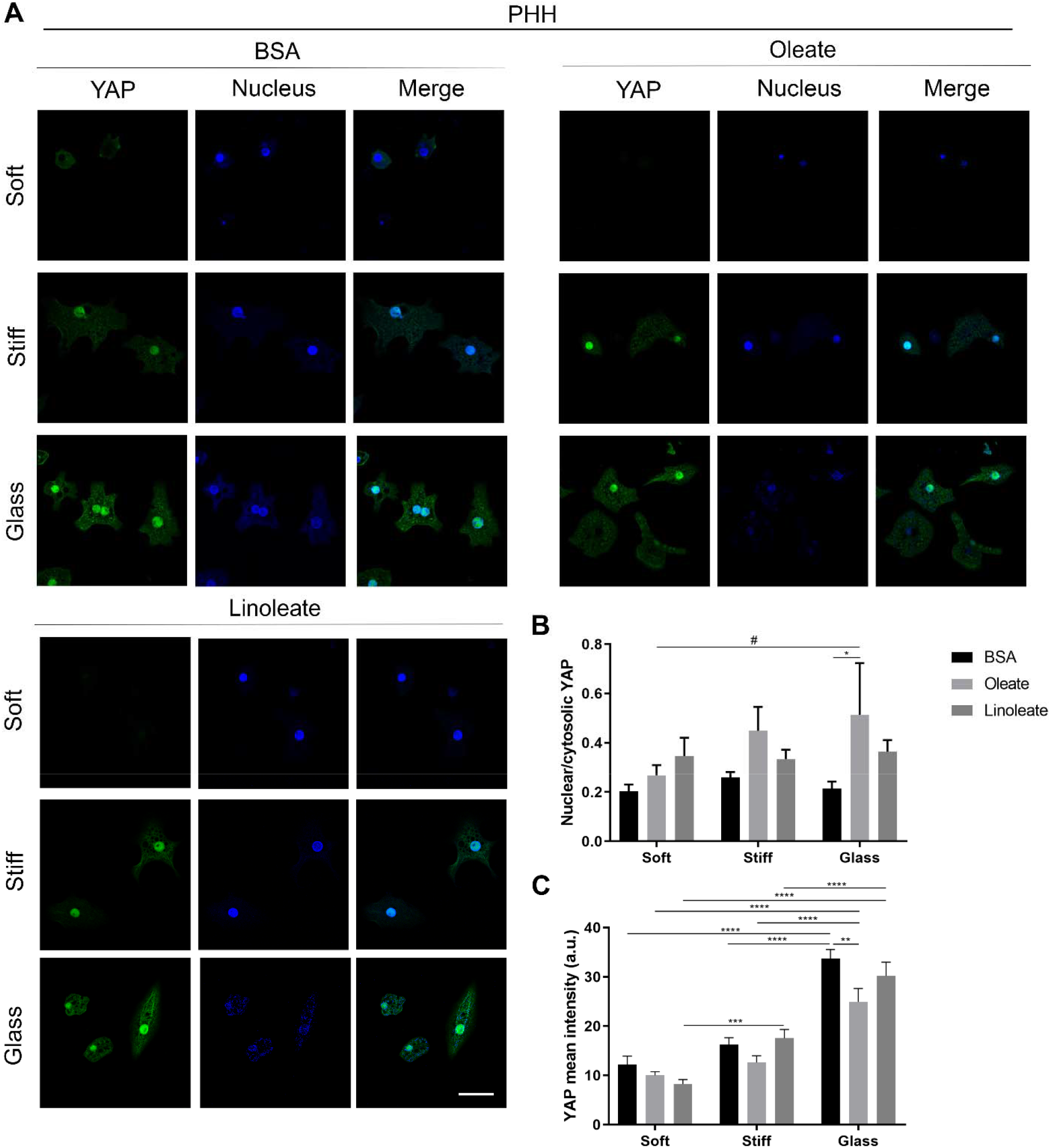
Cells with small lipid droplets show no change in YAP nuclear translocation or mean intensity. **A)** Representative confocal images of PHH seeded on 500 Pa (soft) or 10k Pa (stiff) PAA-collagen gels, or collagen-coated glass, treated with 400 μM oleate or linoleate for 48 h, and stained for YAP (green) and nuclei (blue). **B)** Nuclear to cytosolic YAP ratio was quantified. **C)** YAP mean intensity of the entire cell was measured. Although YAP intensity exhibited some stiffness dependence, fatty acid treatment had no effect. Scale bar, 50 μm. *p≤0.05, ** p≤0.01, ***p≤0.001, ****p≤0.0001, #0.05<p≤0.100. n=29-53 total cells per condition from three independent experiments. Error bars are ± SEM.

Given our finding in cirrhotic livers from NAFLD patients that macrosteatosis was associated with increased YAP nuclear localization (Fig. 3), we added insulin treatment to our lipid loading protocol to generate large lipid droplets in PHH and then determined whether there was a difference between the impact of large versus small droplets on YAP localization. Oleate treatment alone generally resulted in small lipid droplets, while treatment with insulin plus oleate led to overall larger lipid droplets (Fig. 8A). Immunostaining for YAP showed that cells with large lipid droplets had increased nuclear YAP and nuclear deformation in comparison to small lipid droplets, and this reached statistical significance on glass (Fig. 8B, C). Control experiments showed that treatment with insulin alone did not increase, but instead decreased, nuclear YAP compared to BSA controls when seeded on glass (Supplemental Fig. S5).

**Fig. 8.**
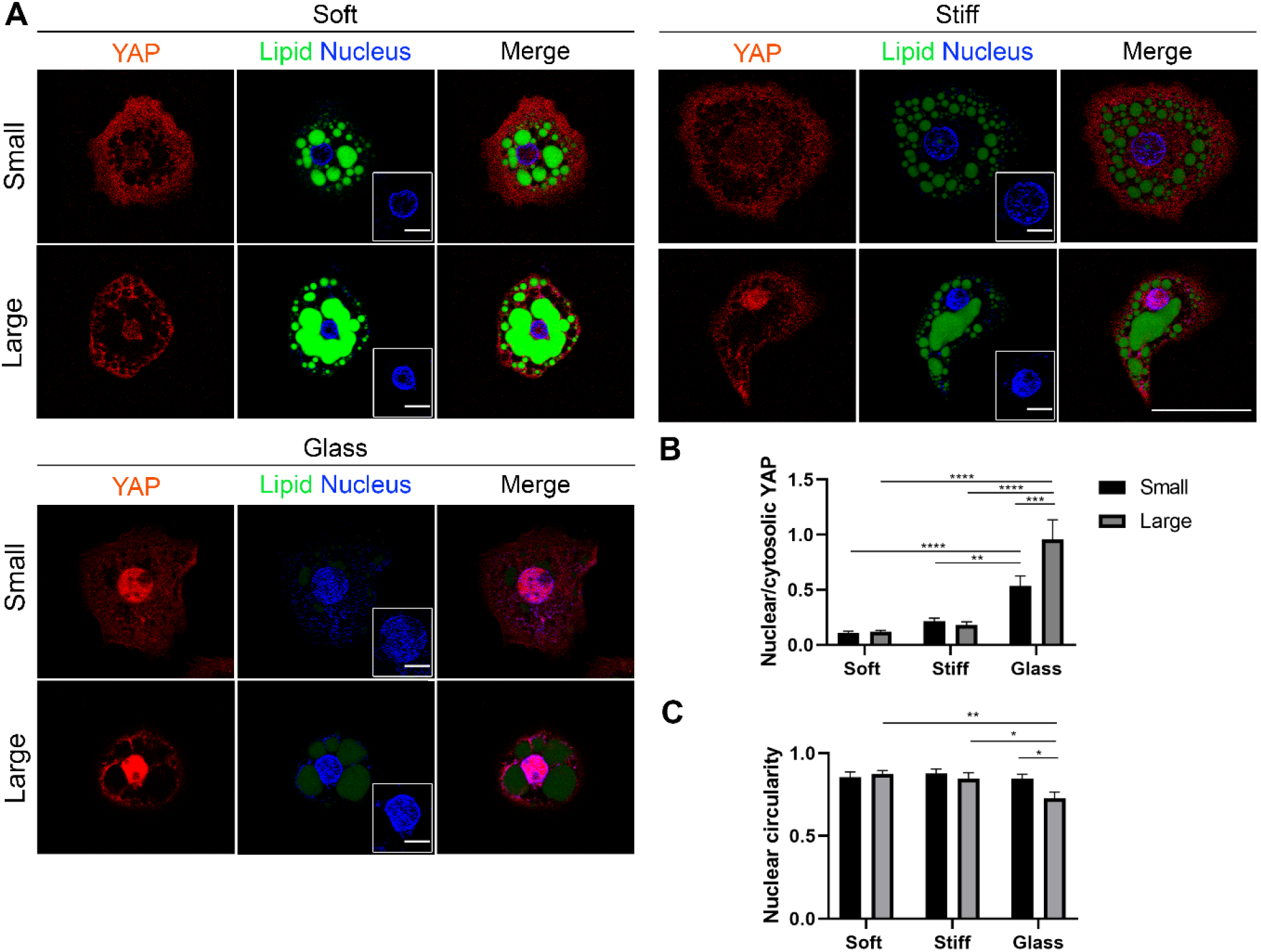
Large lipid droplets are associated with increased nuclear YAP in cells on glass. **A)** Representative confocal images of PHH seeded on 500 Pa (soft) or 10k Pa (stiff) PAA-collagen gels or collagen-coated glass treated with either 400 μM oleate to generate small lipid droplets (microsteatosis) or a combination of 400 μM oleate and 100 nM insulin for large lipid droplets (approaching macrosteatosis). Cells were stained for lipid (green), YAP (red), and nuclei (blue). Intensity was optimized for each stiffness to best visualize YAP staining (A). Nuclei alone are shown in insets. Cells from both treatment conditions were binned into those with predominantly small, medium, or large lipid droplets. **B)** The nuclear to cytosolic YAP ratio and **C)** nuclear circularity were quantified for cells with small versus large droplets. Cells with large lipid droplets had significantly greater nuclear YAP and greater nuclear deformation than those with small lipid droplets when seeded on glass. Inset scale bar, 10 μm. Scale bar, 50 μm. *p≤0.05, **p≤0.01, ***p≤0.001, ****p≤0.0001. n=22-39 total cells per condition from four independent experiments. Error bars are ± SEM.

## DISCUSSION

In this work, we present data suggesting that the presence of lipid droplets alters mechanosensing by hepatocytes. We showed that hepatocytes loaded with lipid droplets had decreased cell spreading as well as disrupted stress fibers and focal adhesions; and that hepatocytes with large, but not small, lipid droplets in both NAFLD cirrhotic livers and PHH in culture had deformed nuclei and increased nuclear localization of YAP. This work underscores the importance of understanding liver mechanical behavior in disease.

Although there are minimal data on the clinical significance of lipid droplet size, some studies suggest an association between macrovesicular steatosis and increased fibrosis, lobular inflammation, hepatocyte damage, insulin resistance, and susceptibility to ischemic injury in comparison to microsteatosis (10, 14, 16, 21, 25). Our *in vivo* data demonstrating an increase in YAP nuclear localization in hepatocytes with large lipid droplets raises the possibility that macrovesicular steatosis serves as a mechanical stress. No other obvious cause of increased mechanical stress in these livers was determined; while cirrhotic livers from patients with NAFLD had more lipid droplets on average than cirrhotic livers from patients without NAFLD, liver stiffness, Sirius Red staining, and content of sGAGs and HA were similar between groups. Stiffness was non-linearly correlated with Sirius Red staining, but not with sGAGs, HA or, notably, lipid content.

In our study, treating hepatocytes in culture with oleate or linoleate alone resulted in the accumulation of small but not large lipid droplets and had no effect on YAP nuclear localization. However, when treated with both oleate and insulin, hepatocytes in culture accumulated large lipid droplets (although not as large as seen in macrosteatosis *in vivo)* and those seeded on glass – a highly stiff substrate – had more nuclear YAP than cells with small lipid droplets.

These findings were consistent with the association we observed between large lipid droplets and YAP localization in cirrhotic livers from patients with NAFLD. Together, our human liver data and fatty acid-loading experiments suggest that lipid droplets, when large enough to displace and deform the nucleus, alter hepatocyte mechanosensing. Moreover, our data corroborate previous work from Elosegui-Artola et al. (9) showing that force transmission to the nucleus, whether by direct application or through contact with a stiff environment, causes stretching of nuclear pores that allows for increased YAP transport to the nucleus. Further understanding of the mechanism by which large lipid droplets drive YAP into the nucleus may provide insight into the development of HCC in soft, non-cirrhotic NAFLD livers (12).

Changes in cell spreading, focal adhesions, and stress fibers were observed after fatty acid-loading, even with the accumulation of only small lipid droplets. This raises the interesting possibility that lipid droplets of varying size may induce different degrees of stress on the nucleus and subsequently different cellular responses. Understanding how these disrupted cytoskeletal elements and YAP nuclear localization affect hepatocyte function will be critical to understanding the relevance of lipid droplet size in disease.

We did not observe significant differences in YAP localization and intensity between soft (500 Pa) and stiff (10k Pa) PAA gels. The dynamic range of hepatocyte mechanosensing on PAA as measured by the extent of cell spreading was determined by Desai et al. to be 75 Pa to 1 kPa (8), and therefore expanding the lower range of gel stiffness might yield more differences in the YAP response. Additionally, our studies were limited to static stiffness. Use of technologies that enable *in situ* substrate stiffening (5) to mimic increasing stiffness of the extracellular matrix as fibrosis progresses should be explored in the future.

Nuclear deformation resulting from large lipid droplets could potentially lead to genetic changes, as observed for cells that migrate through small pores and undergo nuclear deformation that results in genetic instability (11). A large-scale meta-analysis of sequencing data from published work shows that cancers arising from stiff tissues, like muscle and bone, have higher somatic mutation rates than cancers from soft tissues, like marrow and brain (19). Increased collagen within the matrix likely decreases porosity and could lead to genomic mutations in the cells that migrate through these small pores, providing an explanation for how cancers develop in stiff tissues. In the context of NAFLD and HCC, large lipid droplets might induce mechanical force on the nucleus, regardless of the underlying matrix stiffness, and result in cancer-promoting genomic or gene expression changes.

In summary, our *in vitro* experiments indicate that fatty acids prevent hepatocytes from properly sensing or responding to their mechanical environment, as demonstrated by decreased cell spreading, disrupted stress fibers and focal adhesions, and abnormal YAP nuclear localization in the presence of lipid droplets. These droplets may provide a mechanical signal distinct from that exerted by the substrate. This work may provide insight into the mechanisms by which soft NAFLD livers progress to HCC.

## ACKNOWLEDGEMENTS

We are grateful to Emma E. Furth, MD in the Department of Pathology and Laboratory Medicine at the University of Pennsylvania and David E. Kaplan, MD at the Corporal Michael J. Crescenz Veteran’s Administration Medical Center for tissue collection. We acknowledge the Molecular Pathology and Imaging Core of the NIDDK Center for Molecular Studies in Digestive and Liver Diseases (NIH-P30-DK050306) for help in histological processing and the facilities of the Perelman School of Medicine Cell and Developmental Biology Microscopy Core.

## SUPPLEMENTARY MATERIAL

**Supplemental Fig. S1.**
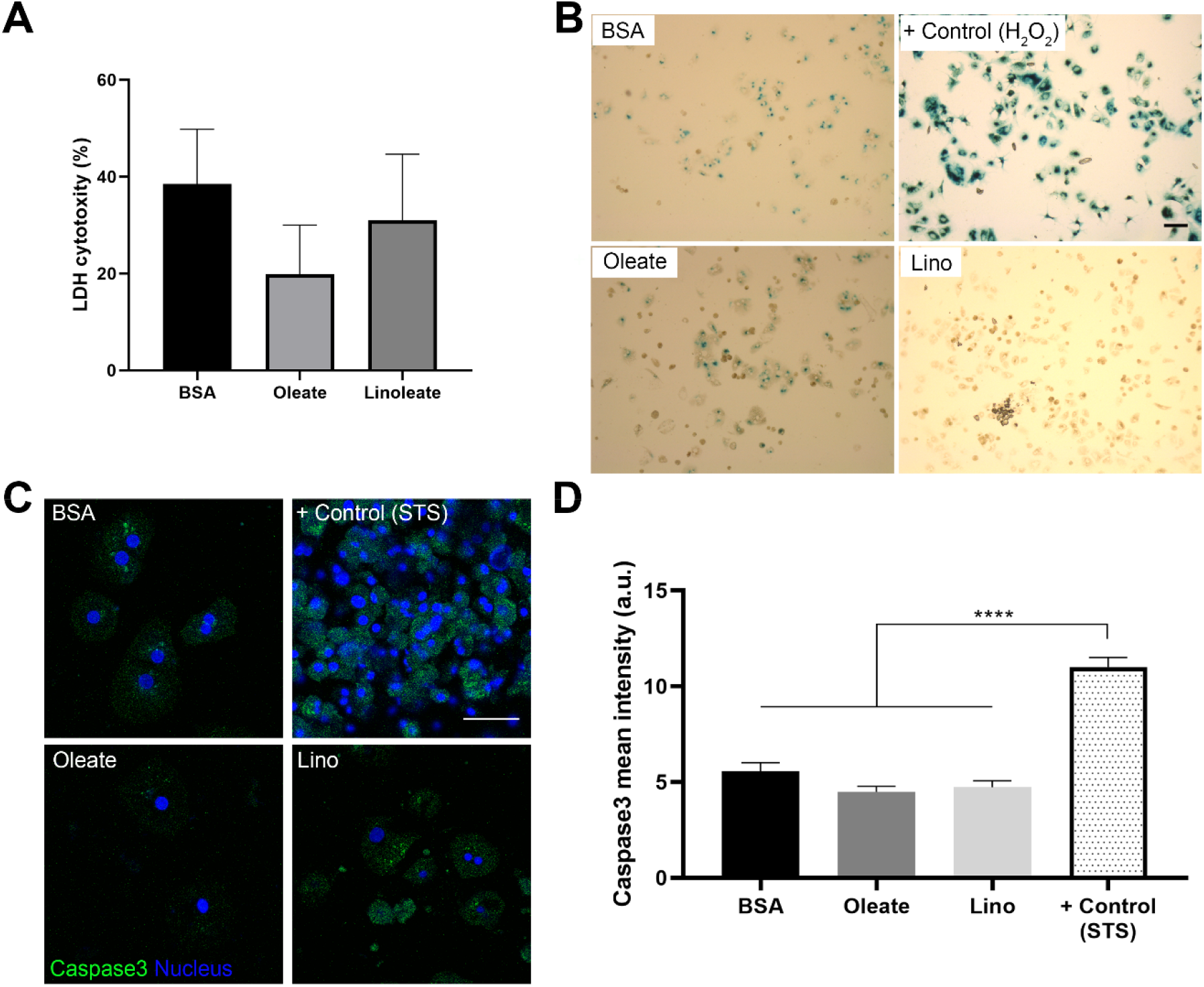
Fatty acid treatment does not cause cytotoxicity or induce senescence or apoptosis. PHH seeded on collagen-coated 96-well plates were treated with 400 μM oleate or linoleate for 48 hours and **A)** assayed for LDH cytotoxicity, **B)** stained for β-galactosidase to assess cell senescence (representative brightfield images), and **C)** stained for caspase3 for apoptosis (representative confocal images). **D)** Caspase3 mean intensity was quantified. Staining of fatty acid-treated cells on PAA gels as opposed to tissue culture plastic similarly showed low intensity caspase3 staining similar to tissue culture plastic (data not shown). Scale bars, 50 μm. ****p≤0.0001. For A, n=3 independent experiments each with triplicate samples. For B, positive control treated with 300 μM H_2_O_2_ for eight days, n=2 independent experiments. For C and D, positive control treated with 5 μM staurosporine (STS) for 20 hours, n=18-23 total cells from 2 independent experiments. Error bars are ± SEM.

**Supplemental Fig. S2.**
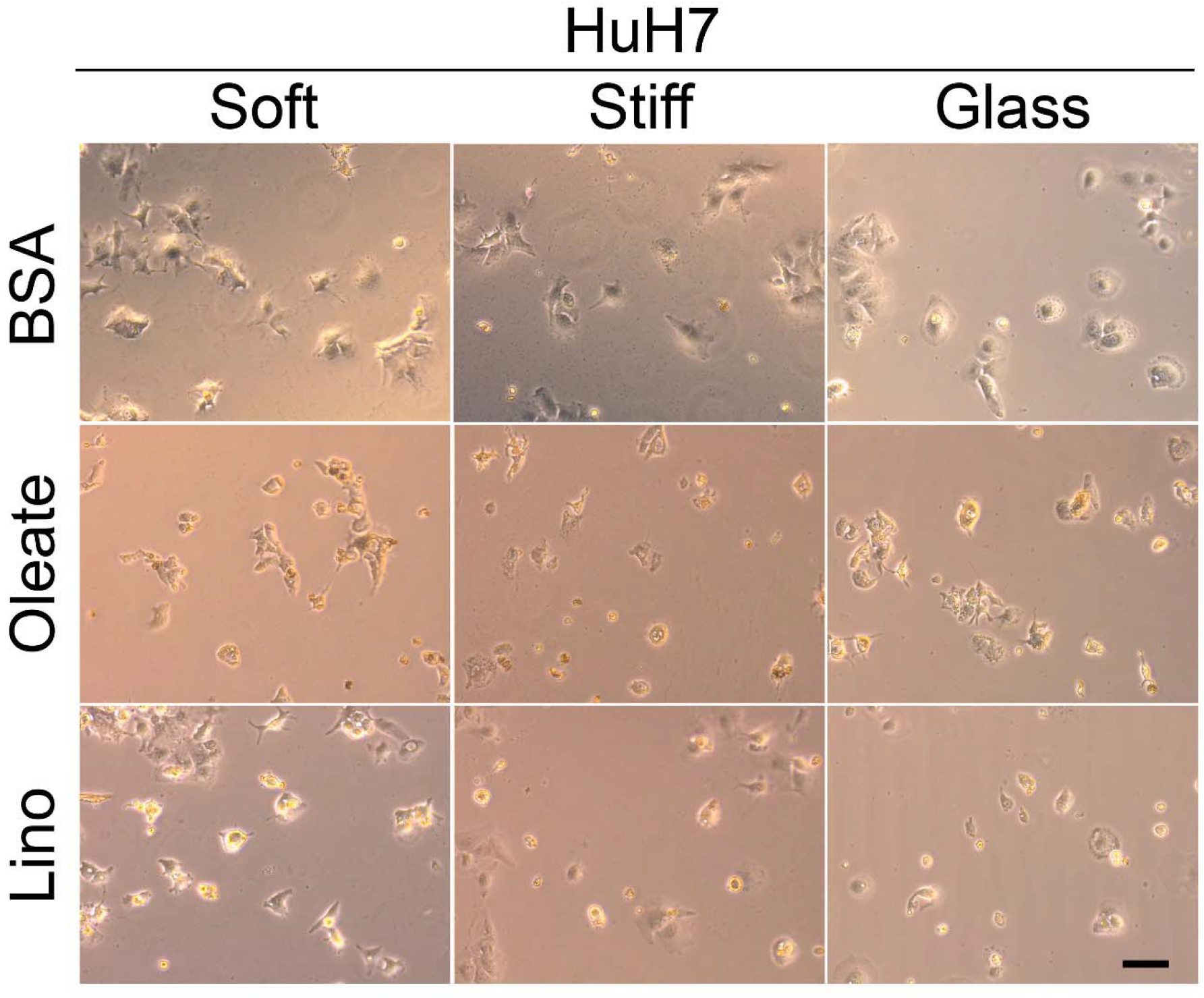
Stiffness-dependent cell spreading decreases in fatty acid-loaded HuH7 cells. Representative live-cell brightfield images of HuH7 cells seeded on 500 Pa (soft) or 10k Pa (stiff) PAA-collagen gels, or glass; treated with 400 μM oleate or linoleate for 48 h. Quantification of cell area, circularity, and solidity are shown in Fig. 5D, E, F. Scale bar, 50 μm. n=60 total cells per condition from three independent experiments.

**Supplemental Fig. S3.**
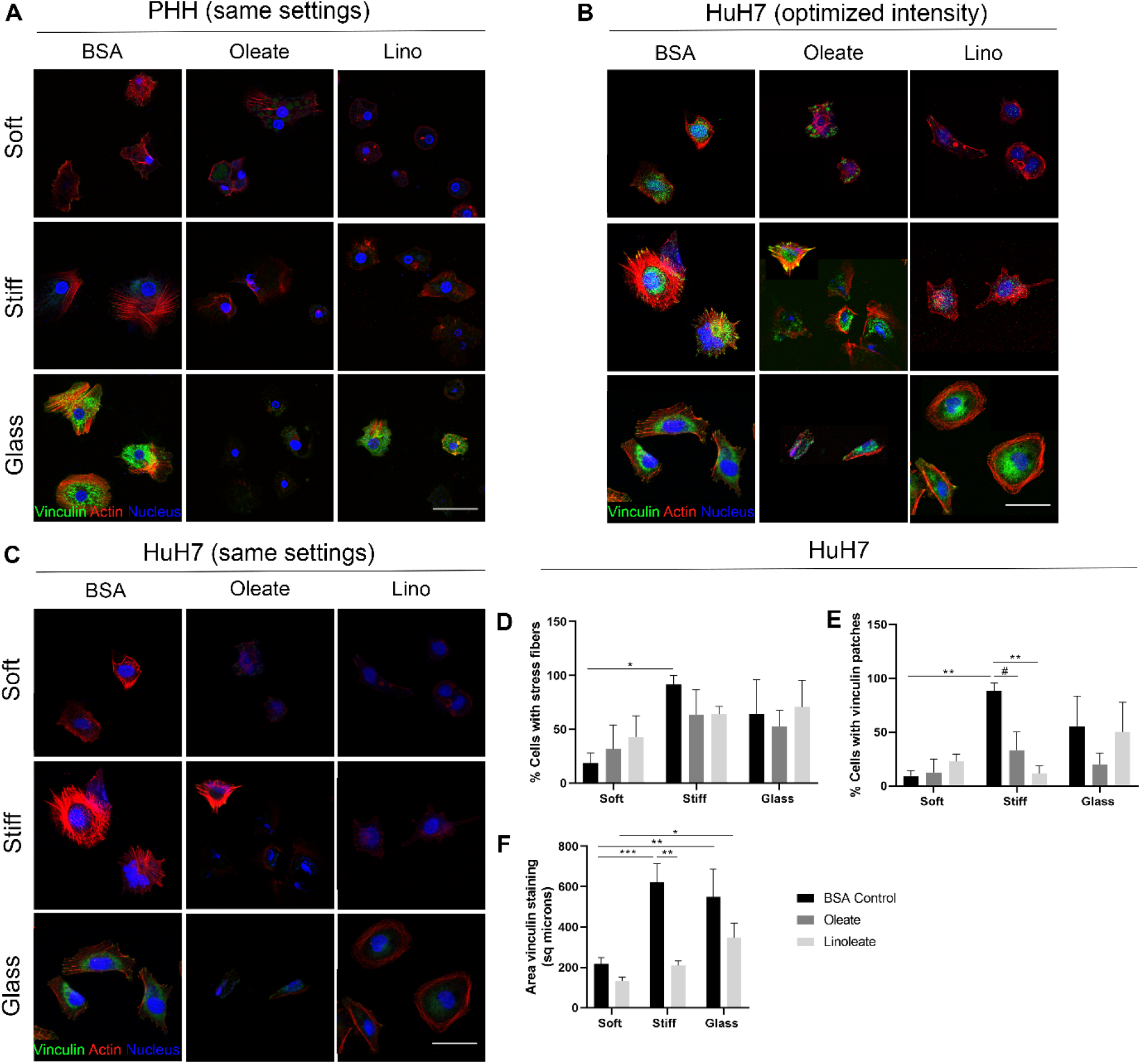
Fatty acid treatment disrupts stress fibers and focal adhesions in PHH and HuH7. Representative confocal images of **A)** PHH and **B and C)** HuH7 cells seeded on 500 Pa (soft), 10k Pa (stiff) PAA-collagen gels, or glass (collagen-coated for PHH), treated with 400 μM oleate or linoleate for 48 h, and stained for vinculin (green), actin (red), and nuclei (blue). Note that panel A is the same as that shown in Figure 6A, but here, imaging settings are the same for all. For Huh7, intensity was optimized for each image to best visualize the vinculin and actin staining **(B)**; vinculin staining was quantified from the original images all taken at the same settings as shown in C. **D)** The percentage of HuH7 cells that exhibited stress fibers, **E)** vinculin patches, as defined by punctate staining located at the end of stress fibers, and **F)** total cell area of vinculin staining (not quantified for oleate-treated cells due to non-specific staining of the lipid droplets). Scale bar, 50 μm. *p≤0.05, ** p≤0.01, ***p≤0.001, ****p≤0.0001, #0.05<p≤0.100. Percent stress fibers or vinculin, n=3 independent experiments. Vinculin area, n=19-35 total cells per condition from three independent experiments. Error bars are ± SEM.

**Supplemental Fig. S4.**
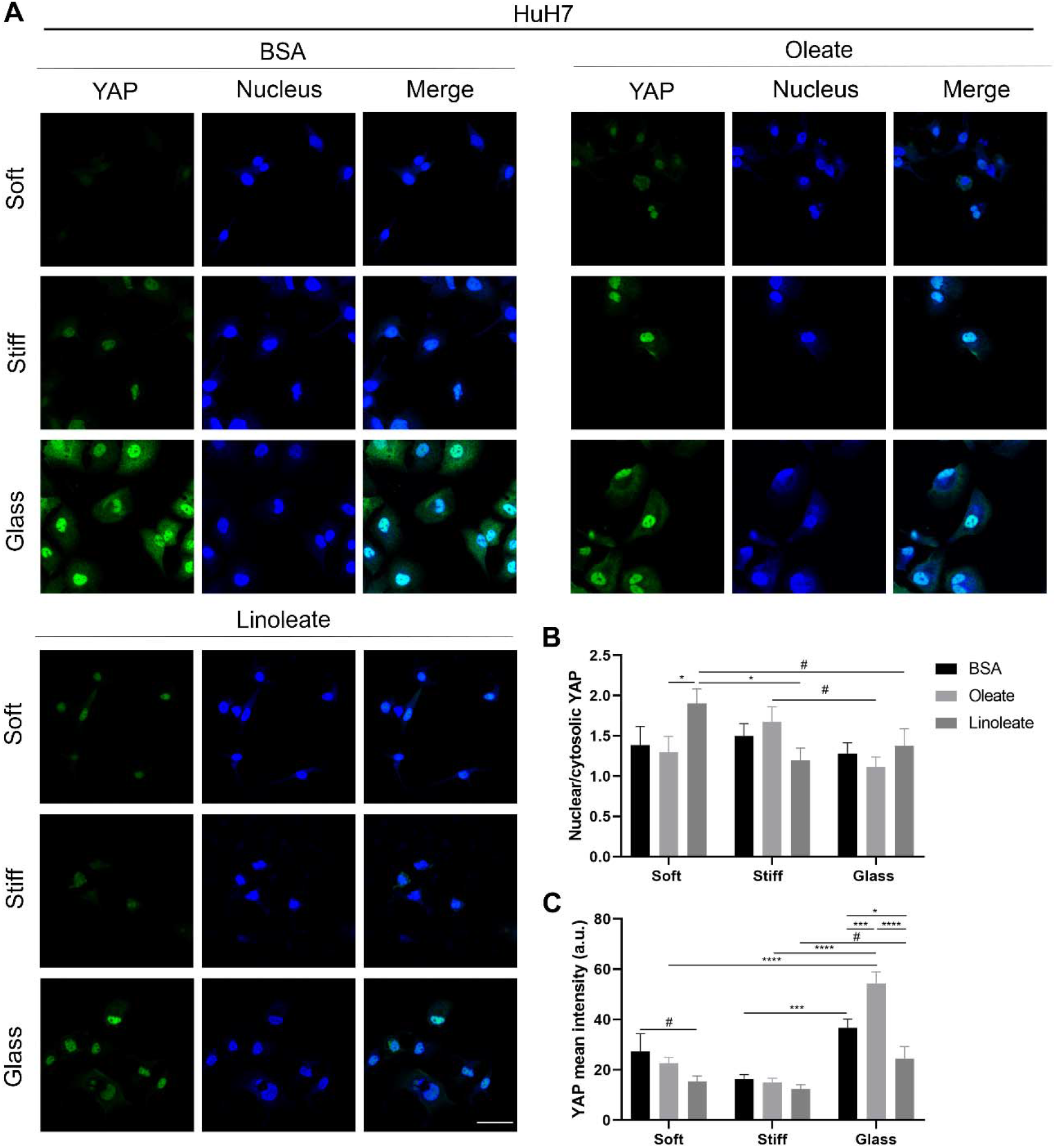
Cells with small lipid droplets show no change in YAP nuclear translocation or mean intensity in Huh7 cells. **A)** Representative confocal images of HuH7 cells seeded on 500 Pa (soft) or 10k Pa (stiff) PAA-collagen gels or glass, treated with 400 μM oleate or linoleate for 48 h, and stained for YAP (green) and nuclei (blue). **B)** Nuclear to cytosolic YAP ratio was quantified. **C)** YAP mean intensity over the entire cell was measured. YAP mean intensity increased on glass, but fatty acid treatment had no effect. Scale bar, 50 μm. *p≤0.05, ***p≤0.001, ****p≤0.0001, #0.05<p≤0.100. n=20-38 total cells per condition from two independent experiments. Error bars are ± SEM.

**Supplemental Fig. S5.**
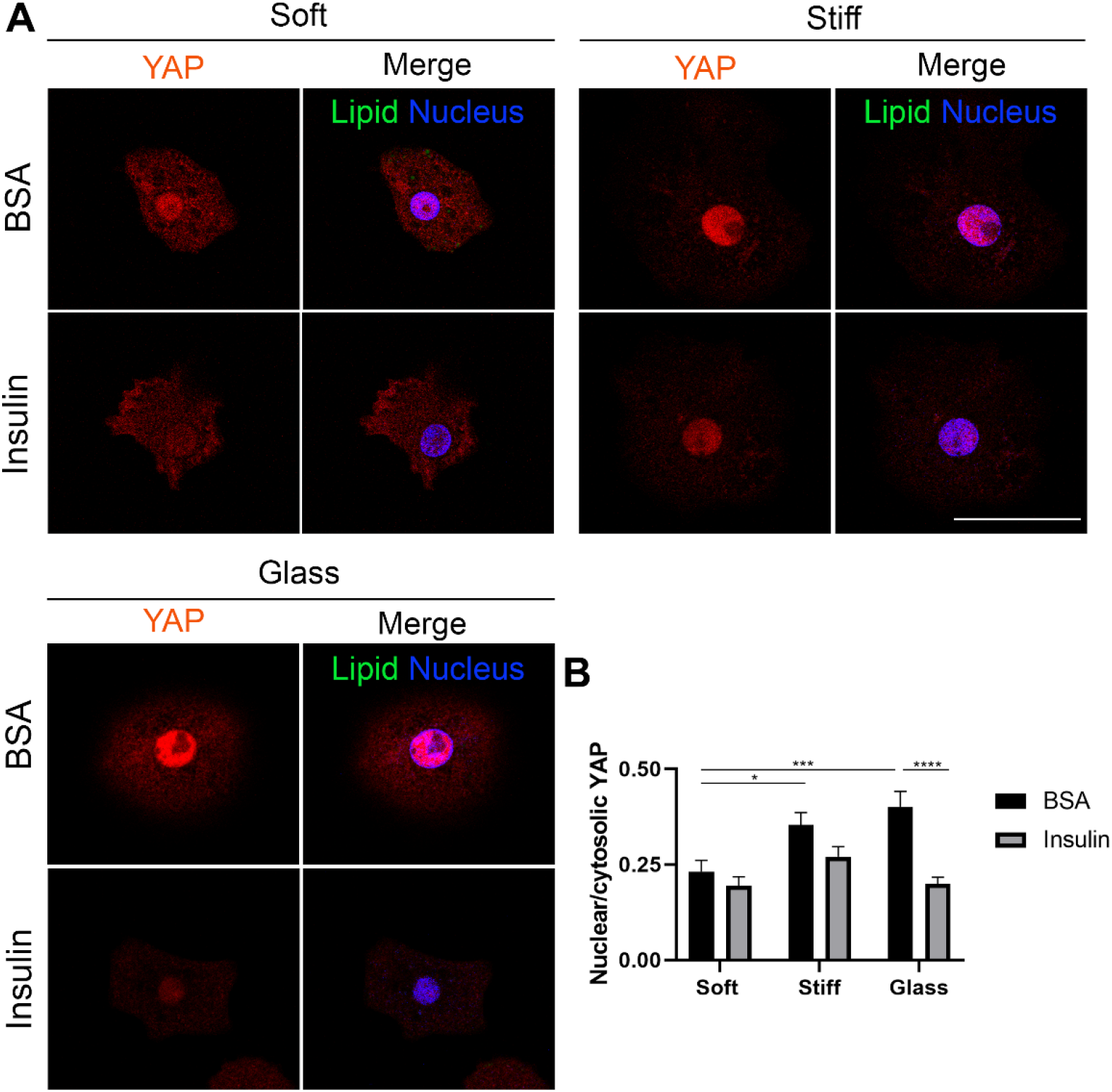
Insulin treatment does not increase nuclear YAP in cells on glass. **A)** Representative confocal images of PHH seeded on 500 Pa (soft) or 10k Pa (stiff) PAA-collagen gels or collagen-coated glass treated with or without 100 nM insulin. Cells were stained for lipid (green), YAP (red), and nuclei (blue). Intensity was optimized for each stiffness to best visualize YAP staining (A). **B)** The nuclear to cytosolic YAP ratio was quantified, and cells treated with insulin had significantly less, but not more, nuclear YAP than BSA controls when seeded on glass. Scale bar, 50 μm. *p≤0.05, ***p≤0.001, ****p≤0.0001. n=34-49 total cells per condition from four independent experiments. Error bars are ± SEM.

## REFERENCES

1. Adler M, Larocca L, Trovato FM, Marcinkowski H, Pasha Y, Taylor-Robinson SD. Evaluating the risk of hepatocellular carcinoma in patients with prominently elevated liver stiffness measurements by FibroScan: a multicentre study. HPB□: the official journal of the International Hepato Pancreato Biliary Association 18: 678–83, 2016.

2. Baker A-M, Bird D, Lang G, Cox TR, Erler JT. Lysyl oxidase enzymatic function increases stiffness to drive colorectal cancer progression through FAK. Oncogene 32: 1863–8, 2013.

3. Baker BM, Trappmann B, Wang WY, Sakar MS, Kim IL, Shenoy VB, Burdick JA, Chen CS. Cell-mediated fibre recruitment drives extracellular matrix mechanosensing in engineered fibrillar microenvironments. Nature Materials 14: 1262–1268, 2015.

4. Boyd NF, Li Q, Melnichouk O, Huszti E, Martin LJ, Gunasekara A, Mawdsley G, Yaffe MJ, Minkin S. Evidence that breast tissue stiffness is associated with risk of breast cancer. PloS one 9: e100937, 2014.

5. Caliari SR, Perepelyuk M, Cosgrove BD, Tsai SJ, Lee GY, Mauck RL, Wells RG, Burdick JA. Stiffening hydrogels for investigating the dynamics of hepatic stellate cell mechanotransduction during myofibroblast activation. Scientific reports 6: 21387, 2016.

6. Cholankeril G, Patel R, Khurana S, Satapathy SK. Hepatocellular carcinoma in nonalcoholic steatohepatitis: Current knowledge and implications for management. World journal of hepatology 9: 533–543, 2017.

7. Cousin SP, Hügl SR, Wrede CE, Kajio H, Myers MG, Rhodes CJ. Free fatty acid-induced inhibition of glucose and insulin-like growth factor I-induced deoxyribonucleic acid synthesis in the pancreatic β-cell line INS-1. Endocrinology 142: 229–240, 2001.

8. Desai SS, Tung JC, Zhou VX, Grenert JP, Malato Y, Rezvani M, Español-Suñer R, Willenbring H, Weaver VM, Chang TT. Physiological ranges of matrix rigidity modulate primary mouse hepatocyte function in part through hepatocyte nuclear factor 4 alpha. Hepatology 64: 261–275, 2016.

9. Elosegui-Artola A, Andreu I, Beedle AEM, Lezamiz A, Uroz M, Kosmalska AJ, Oria R, Kechagia JZ, Rico-Lastres P, le Roux AL, Shanahan CM, Trepat X, Navajas D, Garcia-Manyes S, Roca-Cusachs P. Force Triggers YAP Nuclear Entry by Regulating Transport across Nuclear Pores. Cell 171: 1397–1410.e14, 2017.

10. Han S, Kim G, Lee SK, Kwon CHD, Gwak M, Lee S, Ha S, Park CK, Ko JS, Joh J. Comparison of the tolerance of hepatic ischemia/reperfusion injury in living donors: Macrosteatosis versus microsteatosis. Liver Transplantation 20: 775–783, 2014.

11. Irianto J, Xia Y, Pfeifer CR, Athirasala A, Ji J, Alvey C, Tewari M, Bennett RR, Harding SM, Liu AJ, Greenberg RA, Discher DE. DNA Damage Follows Repair Factor Depletion and Portends Genome Variation in Cancer Cells after Pore Migration. Current Biology 27: 210–223, 2017.

12. Kanwal F, Kramer JR, Mapakshi S, Natarajan Y, Chayanupatkul M, Richardson PA, Li L, Desiderio R, Thrift AP, Asch SM, Chu J, El-Serag HB. Risk of Hepatocellular Cancer in Patients With Non-Alcoholic Fatty Liver Disease. Gastroenterology 155: 1828–1837.e2, 2018.

13. Kulik L, El-Serag HB. Epidemiology and Management of Hepatocellular Carcinoma. Gastroenterology 156: 477–491.e1, 2019.

14. Masterton G, C Ngoh LY, Lockman A, Hayes PC, Plevris J. THE SIGNIFICANCE OF FAT DROPLET SIZE AND THE PROGNOSTIC VALUE OF HYALURONIC ACID IN NON-ALCOHOLIC FATTY LIVER DISEASE (NAFLD): A BIOPSY BASED ANALYSIS. Gut (2011). doi: 10.1136/gut.2011.239301.507.

15. Masuzaki R, Tateishi R, Yoshida H, Goto E, Sato T, Ohki T, Imamura J, Goto T, Kanai F, Kato N, Ikeda H, Shiina S, Kawabe T, Omata M. Prospective risk assessment for hepatocellular carcinoma development in patients with chronic hepatitis C by transient elastography. Hepatology (Baltimore, Md) 49: 1954–61, 2009.

16. McCormack L, Dutkowski P, El-Badry AM, Clavien PA. Liver transplantation using fatty livers: Always feasible? Journal of Hepatology 54 Elsevier B.V.: 1055–1062, 2011.

17. Mittal S, El-Serag HB, Sada YH, Kanwal F, Duan Z, Temple S, May SB, Kramer JR, Richardson PA, Davila JA. Hepatocellular Carcinoma in the Absence of Cirrhosis in United States Veterans is Associated With Nonalcoholic Fatty Liver Disease. Clinical gastroenterology and hepatolog□: the official clinical practice journal of the American Gastroenterological Association 14: 124–31.e1, 2016.

18. Ni Y, Zhao L, Yu H, Ma X, Bao Y, Rajani C, Loo LWM, Shvetsov YB, Yu H, Chen T, Zhang Y, Wang C, Hu C, Su M, Xie G, Zhao A, Jia W, Jia W. Circulating Unsaturated Fatty Acids Delineate the Metabolic Status of Obese Individuals. EBioMedicine 2: 1513–22, 2015.

19. Pfeifer CR, Alvey CM, Irianto J, Discher DE. Genome variation across cancers scales with tissue stiffness – an invasion-mutation mechanism and implications for immune cell infiltration. Current opinion in systems biology 2: 103–114, 2017.

20. Poynard T, Vergniol J, Ngo Y, Foucher J, Munteanu M, Merrouche W, Colombo M, Thibault V, Schiff E, Brass CA, Albrecht JK, Rudler M, Deckmyn O, Lebray P, Thabut D, Ratziu V, de Ledinghen V. Staging chronic hepatitis C in seven categories using fibrosis biomarker (FibroTest™) and transient elastography (FibroScan®). Journal of Hepatology 60: 706–714, 2014.

21. Selzner N, Selzner M, Jochum W, Amann-Vesti B, Graf R, Clavien PA. Mouse livers with macrosteatosis are more susceptible to normothermic ischemic injury than those with microsteatosis. Journal of Hepatology 44: 694–701, 2006.

22. Shikama Y, Ishimaru N, Kudo Y, Bando Y, Aki N, Hayashi Y, Funaki M. Effects of free fatty acids on human salivary gland epithelial cells. Journal of dental research 92: 540–6, 2013.

23. Shoham N, Girshovitz P, Katzengold R, Shaked NT, Benayahu D, Gefen A. Adipocyte stiffness increases with accumulation of lipid droplets. Biophysical journal 106: 1421–31, 2014.

24. Starley BQ, Calcagno CJ, Harrison SA. Nonalcoholic fatty liver disease and hepatocellular carcinoma: a weighty connection. Hepatology (Baltimore, Md) 51: 1820–32, 2010.

25. Stöppeler S, Palmes D, Fehr M, Hölzen JP, Zibert A, Siaj R, Schmidt HH-J, Spiegel H-U, Bahde R. Gender and strain-specific differences in the development of steatosis in rats. Laboratory animals 47: 43–52, 2013.

26. Theise ND, Jia J, Sun Y, Wee A, You H. Progression and regression of fibrosis in viral hepatitis in the treatment era: the Beijing classification. Modern Pathology 31 Nature Publishing Group: 1191–1200, 2018.

27. Umesh V, Rape AD, Ulrich TA, Kumar S. Microenvironmental stiffness enhances glioma cell proliferation by stimulating epidermal growth factor receptor signaling. PloS one 9: e101771, 2014.

28. Wang H-M, Hung C-H, Lu S-N, Chen C-H, Lee C-M, Hu T-H, Wang J-H. Liver stiffness measurement as an alternative to fibrotic stage in risk assessment of hepatocellular carcinoma incidence for chronic hepatitis C patients. Liver international: official journal of the International Association for the Study of the Liver 33: 756–61, 2013.

29. Winer JP, Oake S, Janmey PA. Non-linear elasticity of extracellular matrices enables contractile cells to communicate local position and orientation. PloS one 4: e6382, 2009.

30. Wong RJ, Cheung R, Ahmed A. Nonalcoholic steatohepatitis is the most rapidly growing indication for liver transplantation in patients with hepatocellular carcinoma in the U.S. Hepatology 59: 2188–2195, 2014.

31. Xia T, Zhao R, Feng F, Song Y, Zhang Y, Dong L, Lv Y, Yang L. Gene expression profiling of human hepatocytes grown on differing substrate stiffness. Biotechnology letters 40: 809–818, 2018.

32. Xia T, Zhao R, Liu W, Huang Q, Chen P, Waju YN, Al-Ani MK, Lv Y, Yang L. Effect of substrate stiffness on hepatocyte migration and cellular Young’s modulus. Journal of cellular physiology 233: 6996–7006, 2018.

33. Yang JD, Hainaut P, Gores GJ, Amadou A, Plymoth A, Roberts LR. A global view of hepatocellular carcinoma: trends, risk, prevention and management. Nature Reviews Gastroenterology and Hepatology 16 Nature Publishing Group: 589–604, 2019.

34. Yeung T, Georges PC, Flanagan LA, Marg B, Ortiz M, Funaki M, Zahir N, Ming W, Weaver V, Janmey PA. Effects of substrate stiffness on cell morphology, cytoskeletal structure, and adhesion. Cell motility and the cytoskeleton 60: 24–34, 2005.

